# Role of short-chain fatty acids in the gut-brain axis in schizophrenia: contribution to immune activation and pathophysiology in humans and mice

**DOI:** 10.1101/2020.04.11.021915

**Authors:** Feng Zhu, Wei Wang, Qingyan Ma, Zai Yang, Yajuan Fan, Yanmei Ju, Ruijin Guo, Qi Wang, Xin Mu, Bingbing Zhao, Yuan Gao, Xiaoyan He, Fengjie Gao, Li Qian, Ce Chen, Yunchun Chen, Chengge Gao, Xian-cang Ma

## Abstract

**Objective:** Gut microbiota dysbiosis and aberrant gut-brain functional modules including short-chain fatty acid (SCFA) production and long-lasting immune activation (IA) are presented in schizophrenia. Given the key roles of gut microbiota and SCFA in shaping immunity, we propose that dysbiosis-induced SCFA upregulation could contribute to IA and behavioral symptoms in schizophrenia.

**Design:** Gut microbiota, SCFA, and IA biomarkers were compared between schizophrenic patients and healthy controls. The roles of SCFA in schizophrenia-related IA were analyzed in cultured peripheral blood mononuclear cells (PBMCs) and a mouse model of schizophrenia. The effects of SCFAs on schizophrenia-related phenotypes were analyzed in both human and mouse.

**Results:** Both microbial-derived SCFA and SCFA-producing bacteria were elevated in the guts of schizophrenic patients, and this increased SCFA production in gut was associated with IA in schizophrenia. The microbiome signature underpinning schizophrenia-related IA includes increased diversity and increased SCFA-producing bacteria and inflammation-associated bacteria. The impact of SCFAs on immune responses of cultured PBMC depend on the diagnosis and IA status of donors. Small-molecule serum filtrates from immune-activated schizophrenic patients increased the inflammatory response of PBMCs from healthy volunteers, which can be enhanced and attenuated by SCFAs supplementation and inhibition of SCFA signaling, respectively. Chronically elevated SCFAs in adolescence induced neuroinflammation and schizophrenia-like behaviors in adult mice. Moreover, chronically elevated SCFAs in adult mice prenatally exposed to IA potentiated their expression of schizophrenia-like behaviors.

**Conclusion:** microbiota-derived SCFAs are important mediators of dysregulated gut-brain axis and participant in pathogenesis via enhance IA in schizophrenia.

**Summary:** *Significance of this study:*

1. **What is already known about this subject?**
  ➢ Schizophrenia pathogenesis goes beyond the brain since increasing peripheral abnormalities are revealed including gut microbiota dysbiosis, GI dysfunction, and systemic immune activation (IA).
  ➢Systemic IA/inflammation contributes to the neuroinflammation and brain impairment underlying schizophrenia, and adjunctive immunotherapy can improve psychotic symptoms.
  ➢Short-chain fatty acids (SCFA) mediate the microbiota-gut-brain communication and modulate several pathways involved in schizophrenia, including pathways of immunity and neurotransmitters.
2. **What are the new findings?**
  ➢Patients with schizophrenia displayed increased rates of IA and increased SCFA production compared with healthy controls, and increased SCFA is associated with IA in patients.
  ➢A unique microbiota signature including enriched SCFA-producing bacterial species can distinguish patients with IA from other patients and controls.
  ➢Small molecules in the serum of immune-activated patients with schizophrenia enhance LPS-induced immune response of cultured peripheral blood mononuclear cell (PBMCs), which is partially mediated by SCFA signaling.
  ➢SCFA intake upregulates both peripheral and brain inflammation and potentiates the expression of schizophrenia-like behaviors in mice prenatally exposed to IA.
3. How might it impact on clinical practice in the foreseeable future?
  ➢Interference of SCFA signaling or targeted destruction of SCFA-producing bacteria may provide a new approach for the prevention and treatment of schizophrenia.
  ➢Immune activation status of patients should be an important condition considered when selecting immunotherapy for future precision psychiatric therapy.

## INTRODUCTION

Schizophrenia is one of the most mysterious and devastating mental disorders[1]; however, much of the underlying pathogenic mechanisms remain unknown. The immune/inflammation hypothesis of schizophrenia, based on widely reported disturbances of immune/inflammatory mediators in patients and animal models, posits that dysregulated immune processes interact with the neural system to promote pathogenesis[2]. Increasing evidence indicates that abnormal immune responses contribute to the ongoing pathophysiology of schizophrenia rather than merely the outcome of disease development. Genetic variations that have the strongest associations with schizophrenia risk span across the major histocompatibility complex (MHC) region on chromosome 6.22 that plays a crucial role in immune function[3]. A functional link between immune dysfunction and development of schizophrenia is also suggested by the fact that immune activation is evident for several years prior to the first onset of schizophrenic symptoms[4, 5]. Moreover, nonsteroidal anti-inflammatory drugs and monoclonal antibodies targeting specific inflammatory cytokines or cytokine receptors have shown efficacy at reducing psychopathology in schizophrenic patients[6, 7]. It has been proposed that altered immune pathways partly mediate schizophrenia symptoms, especially impaired cognition[8]. Although great insights have been obtained into the immunological alterations in schizophrenia (see review [9, 10]), the origins of the immune activation and how it is maintained long-term are poorly understood.

The key roles of gut microbiota in shaping the immune system have been demonstrated by numerous studies[11, 12, 13]. Gut microbiota provide tonic immune stimulatory signals to prime the peripheral innate immune system and induce elevated levels of inflammatory cytokines[13]. The interaction between gut microbiota and the host immune system is mainly mediated by microbiota-produced small diffusible metabolites[14]. Short chain fatty acids (SCFAs), the most abundant microbial metabolites in the intestine, play crucial roles in both immune regulation[15, 16, 17, 18, 19] and microbiota-gut-brain communication[20]. Interestingly, many recent studies identified gut microbiota dysbiosis in schizophrenia[21, 22, 23, 24]. Our previous study indicates that the same gut microbiota alterations seen in schizophrenic patients can cause schizophrenia-like abnormal behaviors in mice[25]. Moreover, our recent metagenome-wide association study (MWAS) further identified 83 schizophrenia-associated bacterial species and 27 altered neuroactive potentials of gut microbiota in schizophrenic patients, including SCFA production (manuscript recently accepted by *Nature Communications*). Several bacterial species that are significantly enriched in the gut of schizophrenic patients are SCFA-producing bacteria, including *Bifidobacterium angulatum, Clostridiales bacterium*, and *Lachnospiraceae bacterium*. Neuroinflammation, a common brain abnormality in schizophrenia, is promoted by SCFAs in a mouse model of Parkinson’s disease[26]. Therefore, it is logical to inquire whether SCFA production is dysregulated and can link gut microbiota dysbiosis to aberrant immune activation in schizophrenia.

Potential roles of SCFAs in the pathophysiology of schizophrenia are also likely since they are known to affect metabolic pathways[27, 28], oxidative stress pathways[29, 30], and neurotransmitter pathways[31, 32, 33], all of which are involved in schizophrenia pathology [34]. SCFAs cross the blood brain barrier (BBB) and are internalized by nerve cells to provide energy for these cells, particularly during postnatal brain development[27, 28]. After entering the brain, SCFAs affect cellular oxidative stress functions[29, 30], lipid metabolism[35, 36], regulate the release and synthesis of neurotransmitters[31, 32, 33] and alter microglial homeostasis[37]. Moreover, overstimulation of propionic acid in the brain produces neurocognitive defects, abnormal motor movements, and impaired social interactions in rodents[36, 38, 39]. Therefore, it is important to investigate the role of gut microbiota-SCFA-immune axis interaction in the pathogenesis of schizophrenia, especially schizophrenia-associated immune activation.

## METHODs

### Study design and procedure

Figure 1 shows study design and workflow. Detailed criteria and procedures for recruitment of participants, metagenomic sequencing and bioinformatics, behavioral testing, specimen collection and preparation, and tissues and cell analysis are described in online supplementary methods.

**Figure 1.**
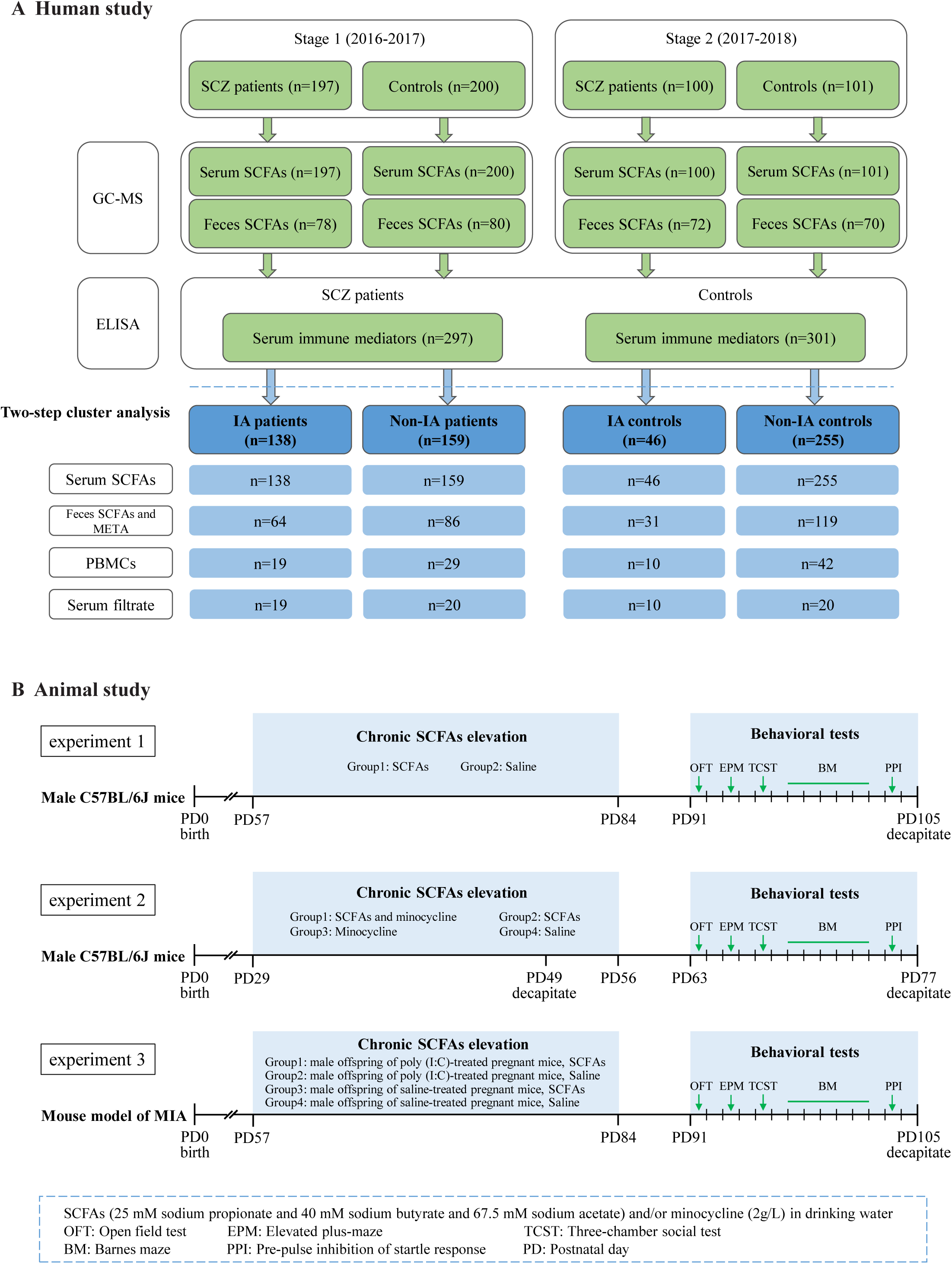
Overview of workflow for the study design. (A) A two-stage of participant recruitment in five hospitals in Shaanxi Province, China. Short-chain fatty acids (SCFAs; acetate, propionate, butyrate) and four immune activation mediators were measured in serum. Patients with schizophrenia (SZ) and healthy controls are classified into four subgroups: immune-activated (IA) patients, non-IA patients, IA controls and non-IA controls. SCFAs in serum and feces, immune response of peripheral blood mononuclear cells (PBMCs) and the immune-activating potentials of small molecule serum filtrates were compared among four subgroups. (B) Two experiments in mice study were conducted to explore excessive SCFA intake in peri-puberty and in adulthood on inflammation level and animal behaviors. Another experiment evaluate the impacts of excessive SCFA intake on behavior symptoms and immune system in mouse model of maternal immune activation (MIA).

### Human study

We explored the relationships among gut microbial-derived SCFAs, gut microbiome, immune functionality, and the diagnosis and phenotypes of schizophrenia. A two-stage cross-sectional study was carried out to compare SCFA concentration between patients with schizophrenia and healthy controls. Systemic immune activation was evaluated for each participant via quantifying four immune mediators in serum, including two markers for monocyte activation: interleukin-6 (IL-6) and soluble CD14 (sCD14), a surrogate marker for T cell activation: soluble interleukin-2 receptor (sIL-2R), and an acute-phase protein (high-sensitive C-reactive protein, hsCRP) [2, 4, 40]. Next, a two-step cluster analysis[41] based on these four immune mediators was applied to classify each participant into one of two distinct categories: “immune-activated” or “non-immune-activated” (see Results). Next, schizophrenia-related phenotypes, SCFA level, structural and functional characteristics of the microbiome, and immune functionality of peripheral blood mononuclear cells (PBMCs) were compared among four subgroups of people, i.e. immune-activated patients or controls and non-immune activated patients or controls. Finally, functional roles of SCFA signaling in immune activation in schizophrenia were analyzed in *in vitro* cultured PBMCs.

### Animal study

The roles of microbial-derived SCFAs in schizophrenia pathology were explored in mice. Three experiments were conducted (see below). Experiments 1 and 2 investigated the impacts of excessive SCFA intake in peri-puberty or adulthood on the expression of schizophrenia-relevant behaviors and immune activation in normal mice. Experiment 3 explored the impacts of SCFA elevation on the expression of schizophrenia-relevant behaviors in adulthood and immune activation in a mouse model of schizophrenia: maternal immune activation (MIA)[42]. The potential roles of inflammation pathways in the effects of SCFAs on mice were evaluated by feeding the anti-inflammatory drug minocycline concurrently with SCFAs.

### Subjects

Participants were recruited as described in our previous studies[25, 43]. This study is a subproject affiliated with a publicly registered clinical trial for gut microbiome investigation in schizophrenia (Identifier: NCT02708316; https://clinicaltrials.gov) that was approved by the Medical Ethics Committee of The First Affiliated Hospital of Xi’an Jiaotong University. All participants signed a consent form. No individuals in this study had physical illness or evidence of infection (all subjects tested normal for hepatic function, renal function, blood-rt (routine test), urine-rt, stool-rt and negative for hepatitis B and C, HIV, HTLV, CMV, syphilis, Chagas disease, and malaria). All participants were physically asymptomatic for gastrointestinal distress.

### Human specimen collection and preparation

Participants had a fasting blood draw at 07:00-09:00 hours. Serum isolated from blood was stored immediately at −80 °C for immunoassay or was further isolated to get small molecule filtrates by centrifugal filtration within three hours after blood draw. The sterile heat-treated serum was centrifuged at 15,000×g for 30 min at 4 °C, and the supernatant was sterile-filtered through a 5-kDa molecular weight cut-off filter (Sartorius Vivaspin® 2 ultrafiltration centrifugal concentrator) to obtain small-molecule serum filtrate. PBMCs were isolated from blood using Ficoll Density Gradient (GE Healthcare, USA) and cultured within two hours of blood draw. Fresh stool sample was collected in hospital and an aliquot was stored in commercial fecal sample collection tubes with *N*-octylpyridinium bromide-based-based reagent (MGIEasy, MGI, Shenzhen, China) for metagenomic sequencing, and a second aliquot was immediately stored at −80 °C for each subject. DNA was extracted from feces as previously described[25].

### Shotgun metagenomic sequencing and MWAS

Metagenomic sequencing was performed on fecal samples from 150 patients and 150 controls. Among them, 39 samples were sequenced in this study and other 261 samples had been sequenced and reported in our previous MWAS for schizophrenia [add ref. after it is online]. The sequencing method, bioinformatic analysis and MWAS were performed as published previously [add ref. after it is online] and as further described in online supplementary methods. Sequence data summary is shown in online supplementary table S3.

### In vitro PBMC study

PBMCs collected from buffy coat were washed with PBS then resuspended in RPMI-1640 medium supplemented with 10% heat-inactivated fetal bovine serum and 1 mM L-glutamine, and 1% Antibiotic-Antimycotic solution. To investigate the effect of SCFAs on immune activation, 2 × 10^6^ cells were treated with 2 mM mixed SCFAs (1.2 mM acetate, 0.4 mM propionate, 0.4 mM butyrate) alone or in combination with GLPG0974 (GLPG, antagonist of GPR43; 0.1 mM) and b-hydroxybutyrate (SHB, antagonist of GPR41; 5 mM) for 16 h and were then exposed to lipopolysaccharide (LPS, 5 µg/mL) for 12 h. To investigate the effects of serum filtrate on immune activation, 2 × 10^6^ PBMCs from healthy volunteers were incubated with PBS or small-molecule serum filtrate alone or in combination with mixed SCFAs, or in combination with GLPG and SHB for 24 h. Supernatants were collected and stored at −80 °C for cytokine measurement. mRNA level of immune-related genes and *GPR41* and *GPR43* were measured via qPCR (supplementary table S2).

### Animal studies

Male C57BL/6J mice were used for all experiments and were housed under standard SPF environment with food and water provided ad libitum. All experiments were approved by the Animal Care and Use Committee of Xi’an Jiaotong University. In experiment 1, mice were fed with SCFAs (25 mM sodium propionate and 40 mM sodium butyrate and 67.5 mM sodium acetate[37]) and minocycline (2g/L)[44], SCFAs alone, minocycline alone in drinking water, or sodium-matched water for 4 weeks, from postnatal days (PD) 29-56, then tested on a series of behavioral tasks one week later. In experiment 2, mice were fed with SCFAs or sodium-matched water during postnatal days (PD) 57 - 84 and underwent behavior evaluation one week later. In experiment 3, mice prenatally exposed to immune stimulation or sodium-matched water were treated with combined SCFAs and minocycline, SCFAs alone, minocycline alone, or saline during PD 57 - 84 and underwent behavioral testing one week later. MIA was established as published previously[42]. Dosage of agents and behavioral tests were identical in all three experiments. Behavioral testing was performed in the following order: open field test (OFT), elevated plus maze (EPM), three-chamber social test (TCST), Barnet maze (BM), and pre-pulse inhibition of startle response (PPI). Mice were rapidly decapitated 24 hours after the final behavioral test. Serum was harvested from the neck, and prefrontal cortex (PFC) and hippocampus were dissected from the brains. TNF-α and IL-6 were measured in serum using ELISA kits.

### SCFA quantification

Acetate, propionate, and butyrate in serum of the discovery set subjects were quantified via gas chromatography–triple quadrupole mass spectrometry (GC-QQQ-MS) assay. Serum SCFAs of the validation set subjects and the mice and all fecal SCFAs were measured via gas chromatography–mass spectrometry (GC-MS). Detailed procedures and parameters for SCFAs quantification in GC-QQQ-MS and GC-MS are shown in online supplementary methods and online supplementary Table S1, respectively.

### Immunoassay

Hs-CRP was measured using immunoturbidimetric assay (TBA-200FR, Tokyo, Japan). Pro-inflammatory cytokines in human serum, supernatant of cultured human PBMCs and mouse colon, serum, and brain were quantified using commercially available enzyme-linked immunosorbent assay kits according to the manufacturer’s instructions.

The identifier, assay sensitivity and intra- and inter-assay coefficients of variation of these kits are shown in online supplementary Table S2. The activity of NF-κB in nuclear extracts of PBMCs was measured using a commercially available NF-κB (p65) Transcription Factor Assay (Cayman Chemicals) following the manufacturer’s instructions.

### Statistical analyses

Continuous variables were expressed as mean ± standard deviation (SD), mean ± standard error of the mean (SEM), or median with interquartile range (IQR). Data were tested for normal distribution using the D’Agostino & Pearson normality test. Parametric (Student’s *t*-test and analyses of variance) or nonparametric group comparisons (Mann-Whitney U and Kruskal-Wallis H-tests) were applied for continuous variables as appropriate. χ^2^ tests (for frequencies) were calculated for the difference in distribution of categorical variables. Spearman rank correlation analyses were performed for SCFA levels, inflammatory factors, gut bacterial abundance, and gut microbiota diversity in both groups. Multivariate general linear model (GLM) analysis was performed to investigate whether SCFAs or immune factors were influenced by diagnosis and immune activation status. Gender, age, BMI, smoking status, and alcohol drinking status were included in the model as potential confounders. When the distribution of a variable was skewed, natural log (Ln) transformations were used before GLM analysis to reduce variance and outlier influence. The potential diagnostic values of SCFAs and gut microbial classifiers were evaluated by a receiver-operating characteristic curve (ROC) analysis with calculation of the value of the area under the ROC curve (AUC), as well as by a stepwise binary logistic regression. Statistical significance was set at *P* < 0.05. Adjusted *P*-values were obtained for multiple comparisons using the Benjamini and Hochberg correction (false discovery rate FDR). All analyses were carried out using the free software package R (http://cran.rproject.org/) and GraphPad Prism 7.0.

## Results

### Upregulation of microbial-derived SCFAs in schizophrenic patients

Demographic characteristics of patients and controls are shown in table 1. In discovery set, acetate, propionate, butyrate, and total concentration of these three types of SCFA (referred to total microbial SCFA thereafter) significantly increased in the patients compared to the controls (*P* < 0.0001 for each comparison; figure 2A-D; supplementary table S4). Multivariate GLM analyses also revealed serum SCFAs were mainly affected by diagnosis (*P* < 0.0, supplementary table S5). After adjusting potential confounders, acetate, propionate, and total microbial-derive SCFA significantly predicted the diagnosis of schizophrenia (all *P* < 0.001; supplementary table S6), and total microbial SCFA had highest accuracy for diagnosis of schizophrenia (*P* < 0.0001, AUC = 0.7542; figure 2E). The elevations of acetate, propionate, butyrate and total microbial-derived SCFA in the serum of the patients compared to those of the controls as well as their associations with the schizophrenia risk were replicated to be also statistically significant in the validation set (100 patients vs. 101 controls; all *P* < 0.01; supplementary Tables 4, 6). Schizophrenic patients displayed poor performance in all cognitive tasks (all *P* < 0.0001; table 1). Moreover, SCFAs significantly negatively correlated with cognitive function after controlling age, gender, education, and BMI, i.e. the patients with higher SCFA level displayed more severe cognitive impairment (supplementary table 7a). Forty-five patients were followed up three months later. Pair-wise comparison of serum SCFA and cognitive function for treatment effect in these patients revealed treatment decreased serum SCFA and improved cognition, accompanied with significant correlations between their changes caused by treatment (all *P* < 0.01; supplementary table 7b).

**Figure 2.**
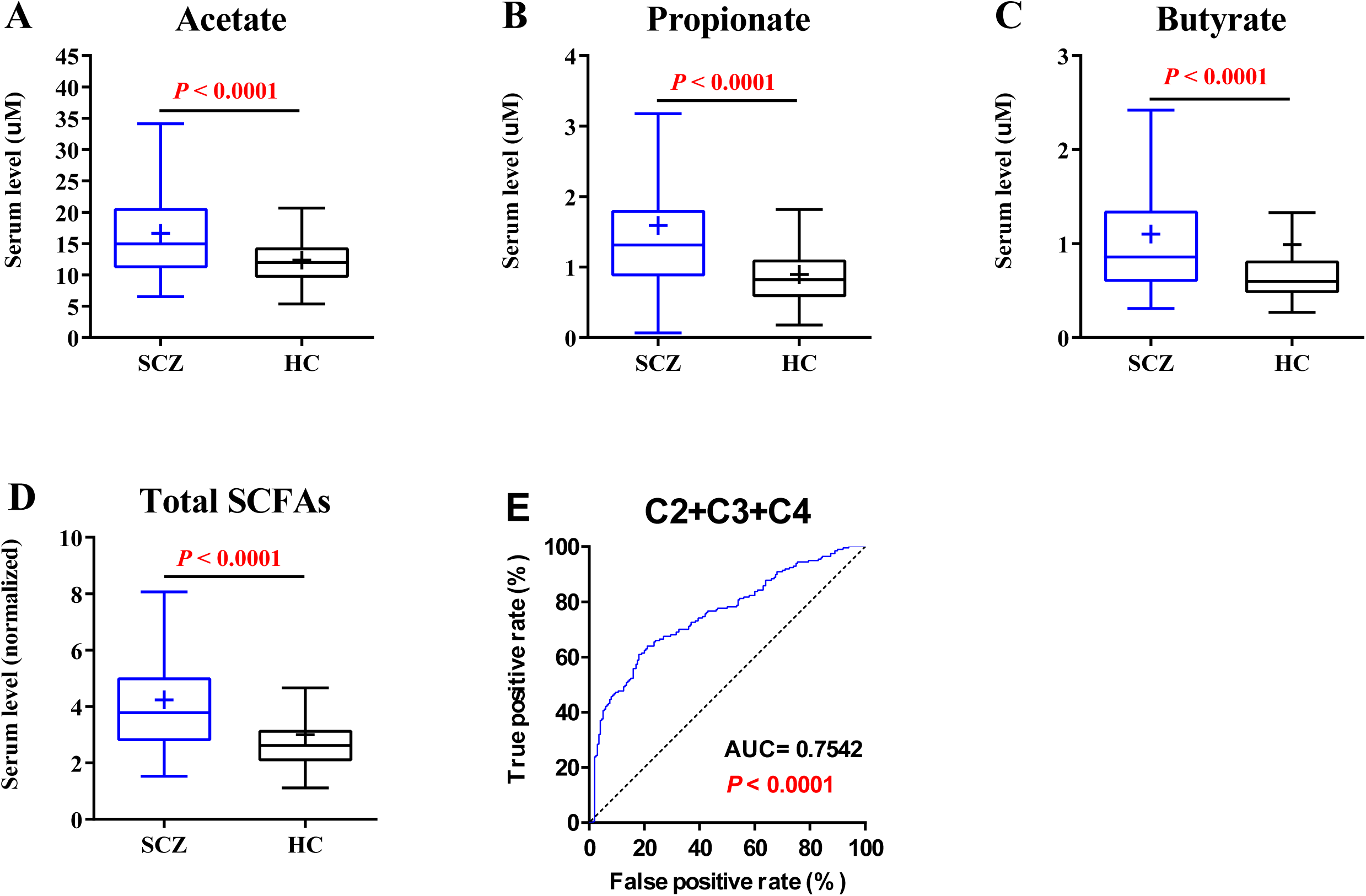
Serum short-chain fatty acids (SCFAs) is differentially distributed between schizophrenic patients and healthy controls and predicts the disease risk. (A) acetate, (B) propionate, (C) butyrate, (D) their total concentration (main microbial-derived SCFAs) in serum were compared between the patients (n = 197) and the controls (n = 200). (A-D) The dot represents one value of serum concentration from an individual participant; The box represents the median and interquartile ranges (IQRs) between the first and third quartiles; whiskers represent the lowest or highest values within 1.5 times IQR from the first or third quartiles; Outliers are not shown; *P* value was determined by Mann Whitney test. (E) Classification performance of total microbial SCFAs in serum to distinguish schizophrenia from controls was assessed by area under the receiving operational curve (AUC).

### Increased SCFA excretion and SCFA-producing bacteria in the gut of schizophrenia patients

To determine the origin of increased serum SCFAs in schizophrenic patients, we subsequently quantified acetate, propionate, and butyrate in the feces of 78 patients and 80 controls who were selected randomly from the discovery set. GLM revealed all three types of SCFAs were increased significantly in the feces of SCZ patients, after adjusting for measured confounders (all *P* < 0.05; figure 3A-C; supplementary table S8a). Moreover, there were significantly correlations between fecal SCFAs and SCFAs in serum in patients (*r* = 0.236∼0.394, all *P* < 0.001), but nor in controls (supplementary table S8b). Increased microbial SCFAs in feces and the positive association between fecal and serum SCFAs were again confirmed by data from 72 patients and 70 controls selected randomly from the validation set (supplementary table S8). Our previous study indicates that some bacterial species that produce SCFAs are enriched in the gut of schizophrenic patients[45]; therefore, we next sought to seek out the bacterial species underpinning the upregulation of serum and fecal SCFAs in schizophrenia. In line with our previous study[45], gut microbiota α-diversity was increased in patients with schizophrenia, and was positively associated with SCFA concentration in serum (Shannon index; *r* = 0.138∼0.204, *P* = 0.016∼0.0002) but not in feces (Supplemental Table S9).

**Figure 3.**
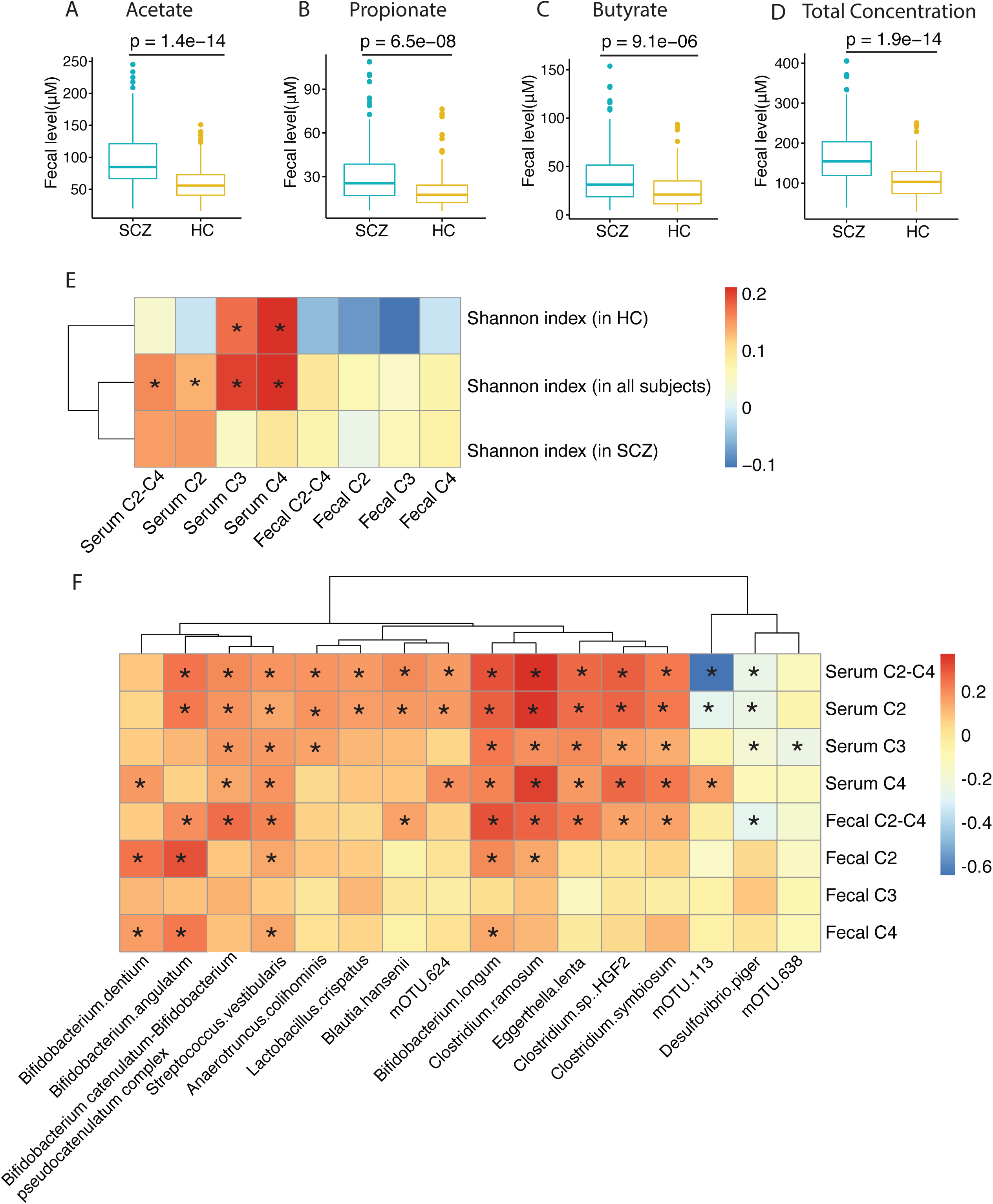
Microbial short-chain fatty acids (SCFAs) and SCFA-producing bacteria were upregulated in the gut of schizophrenia patients and correlated with the serum SCFAs. (A-D) Fecal acetate, propionate, butyrate and the sum of these three types of SCFAS were elevated in the patients with schizophrenia. The box represents the median and interquartile ranges (IQRs) between the first and third quartiles; whiskers represent the lowest or highest values within 1.5 times IQR from the first or third quartiles. *P* value were determined by Mann Whitney test. (E) Spearman correlations of gut microbiota with the SCFAs in feces and with those in serum in the patients with schizophrenia.

Spearman correlation identified 129 species of gut bacteria that correlated with at least one of the serum SCFAs in patients (*P* < 0.05, FDR < 0.1, online supplementary table S10). Of these, 42 showed moderate-to-strong correlations (|rho| > 0.20). Among these 42 mOTUs associated with serum SCFAs in patients, 16 species of gut bacteria were also significantly enriched in gut of patients compared to controls (q < 0.05; online supplementary table S10; figure 3F).

### Higher circulating SCFAs are present in a subgroup of schizophrenic patients with immune activation

To determine the relationship between increased production of SCFAs and abnormal immune activation in schizophrenia, we quantified four biomarkers for immune activation: C-reactive protein, IL-6, sIL-2R, and sCD14 in serum of all participants. All four biomarkers were significantly upregulated in schizophrenic patients compared to controls (all *P* < 0.0001, figure 4A-D). There were small to moderate inter-correlations among these four biomarkers in patients after adjustment for potential confounders (all *P* < 0.001, r: 0.09∼0.54; supplementary table S11). Several significantly positive correlations existed between microbial-derived SCFAs and the four biomarkers of immune activation in the serum of patients (all *P* < 0.0001, supplementary table S11). The strongest correlation was between total microbial-derived SCFAs and IL-6 (*r* = 0.445, *P* < 0.0001; figure 4E). Multivariate GLM analysis showed significant interaction effects of diagnosis and total microbial SCFAs on serum IL-6 and sCD14 (*P* < 0.001, supplementary table S12a).

**Figure 4.**
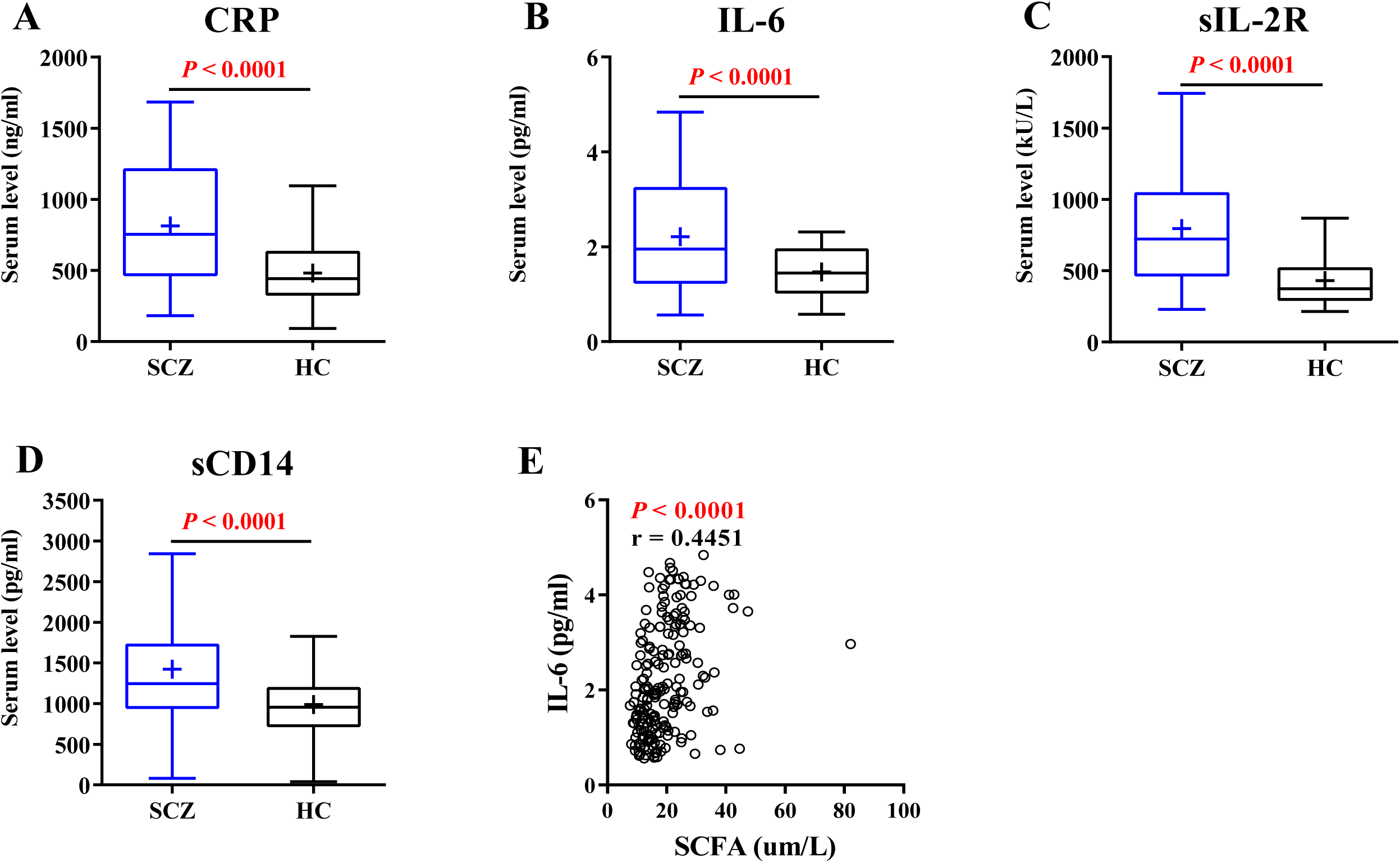
A subgroup of schizophrenic patients characterized by elevated serum biomarkers for immune activation display expressed higher short-chain fatty acids (SCFAs) in serum. (A-D) C-reactive protein, IL-6, sIL-2R, and sCD14 were elevated in serum of the patients with schizophrenia compared with controls. The box represents the median and interquartile ranges (IQRs) between the first and third quartiles; whiskers represent the lowest or highest values within 1.5 times IQR from the first or third quartiles. Outliers are not shown. *P* value were determined by Mann Whitney test. (E) Correlation of four biomarkers of immune activation with SCFAs.

A two-step cluster analysis on these four immune biomarkers identified two distinct clusters; “immune-activated group,” referring to a pattern of relatively higher biomarker levels and “non-immune-activated group” for those with a pattern of relatively decreased biomarker levels. The immune-activated group was characterized by immune biomarker concentrations of >85% subjects in this group had biomarker concentrations higher than *mean + 2 × SD* of the biomarker concentrations in the non-immune-activated group (supplementary table S12b; *P* < 0.0001). The proportion of subjects with immune activation in the patient group was 138/297 and for controls this was 46/301. The proportion of the patients with immune remained similar between the discovery cohort (95/197) and the validation cohort (43/100), suggesting good reproducibility of clustering schizophrenia by these immune biomarkers. No significant differences in most demographic features existed among the four subgroups of subjects (*P* > 0.05), except for significantly greater age (*P* = 0.025) and nominally elevated BMI (*P* = 0.061) in the controls with immune activation than those without (supplementary table S13a). Subgroup comparison indicated that immune-activated patients had higher serum SCFAs and fecal SCFAs and poorer cognition than the patients without immune activation and all controls (all *P* < 0.001, supplementary table S13b).

### Microbiome signature for the immune activation in schizophrenia

Considering key roles of gut microbiota in shaping the host immune system, we hypothesized that dysbiotic microbiota may underpin immune activation in schizophrenia. Therefore, we sought to identify the unique microbial signature correlating with immune activation in schizophrenia via a metagenome-wide association study. Immune-activated patients had greater α-diversity of gut microbiota at the mOTU level than both non-immune-activated patients and controls (Shannon index; figure 5A; online supplementary tables S14). β-diversity based on Bray-Curtis distance also varied among four subgroups of people and non-immune-activated healthy controls had increased Bray-Curtis distance compared other three subgroups (figure 5B). PCoA based on Bray-Curtis dissimilarity of mOTUs showed that the overall faecal microbiota composition was different between immune-activated patients and non-immune-activated patient (figure 5C). Differentially enriched mOTUs in gut of each subgroup are shown in online supplementary table S15. Compared to non-immune activated controls, 20 mOTUs were differentially enriched in the gut of immune-activated patients, 5 mOTUs in non-immune-activated patients, and just one mOTU in immune-activated controls after Bonferroni correction (all *P* < 0.05/360). Top 5 differentially-enriched mOTUs in immune-activated patients compared with non-activated controls includes Eggerthella lenta, Bifidobacterium longum, Clostridium sp. HGF2, Akkermansia muciniphila, Bifidobacterium angulatum (all *P* < 0.00001). Notably, the differentially enriched bacteria in immune-activated patients included several SCFA-producing bacteria (supplementary table S15). Next, we constructed a set of random forest disease classifiers based on gut mOTUs to identify taxa that best distinguish four subgroups of people. A five-fold cross-validation procedure was conducted ten times on 150 patients and 150 controls. We found that three models displayed good efficacy to distinguish immune-activated patients from other three subgroups with an AUC of 0.84∼88 (figures 5D, E, F). We then constructed a mOTU network to depict the cooccurrence correlation between the schizophrenia-associated gut bacteria under the condition of immune activation and non-activation, respectively (figures 5G, H). Immune-activated schizophrenia-enriched mOTUs were more interconnected than control-enriched mOTUs (Spearman’s correlation coefficient <−0.3 or >0.3, *P* < 0.05).

**Figure 5.**
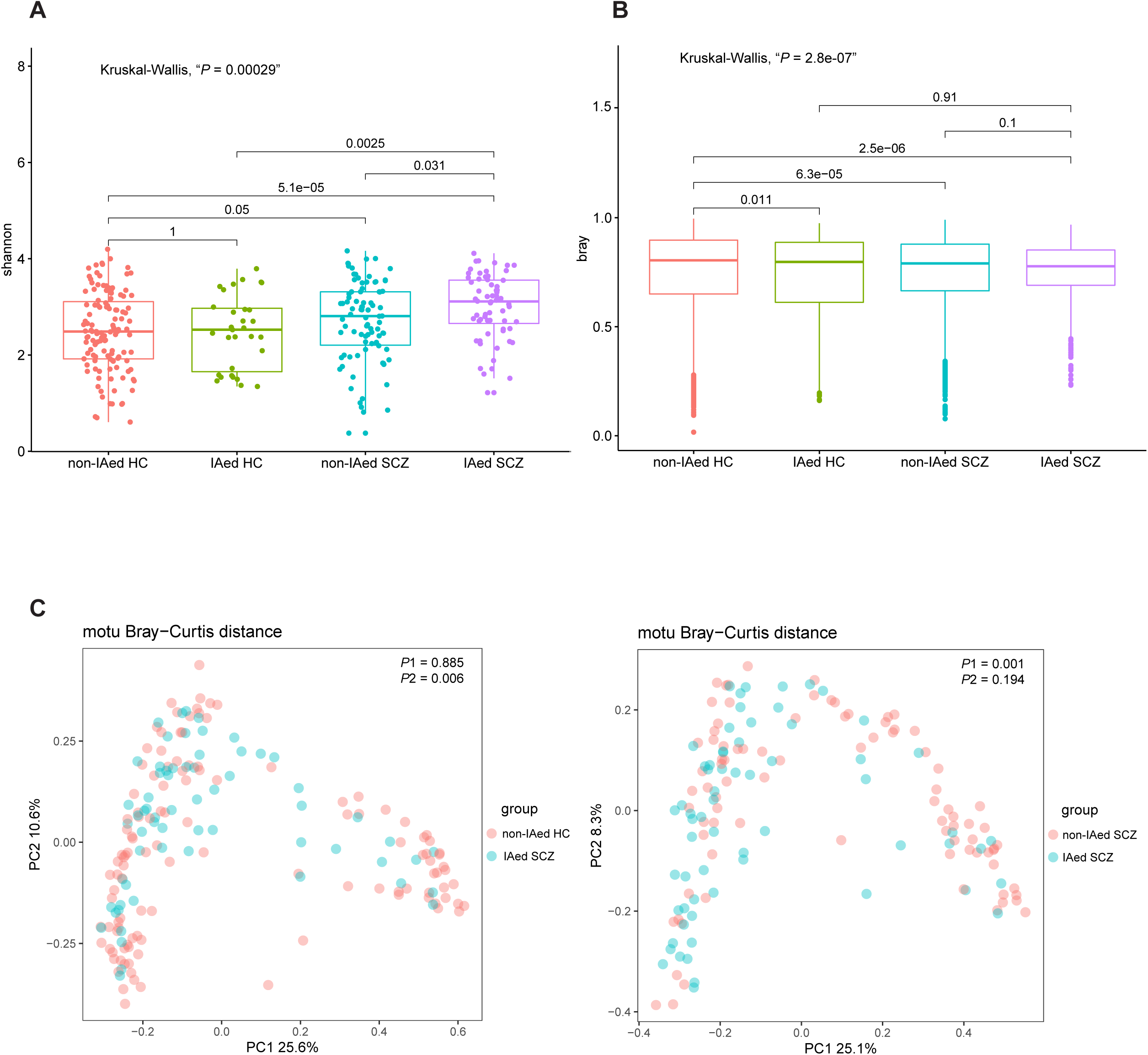
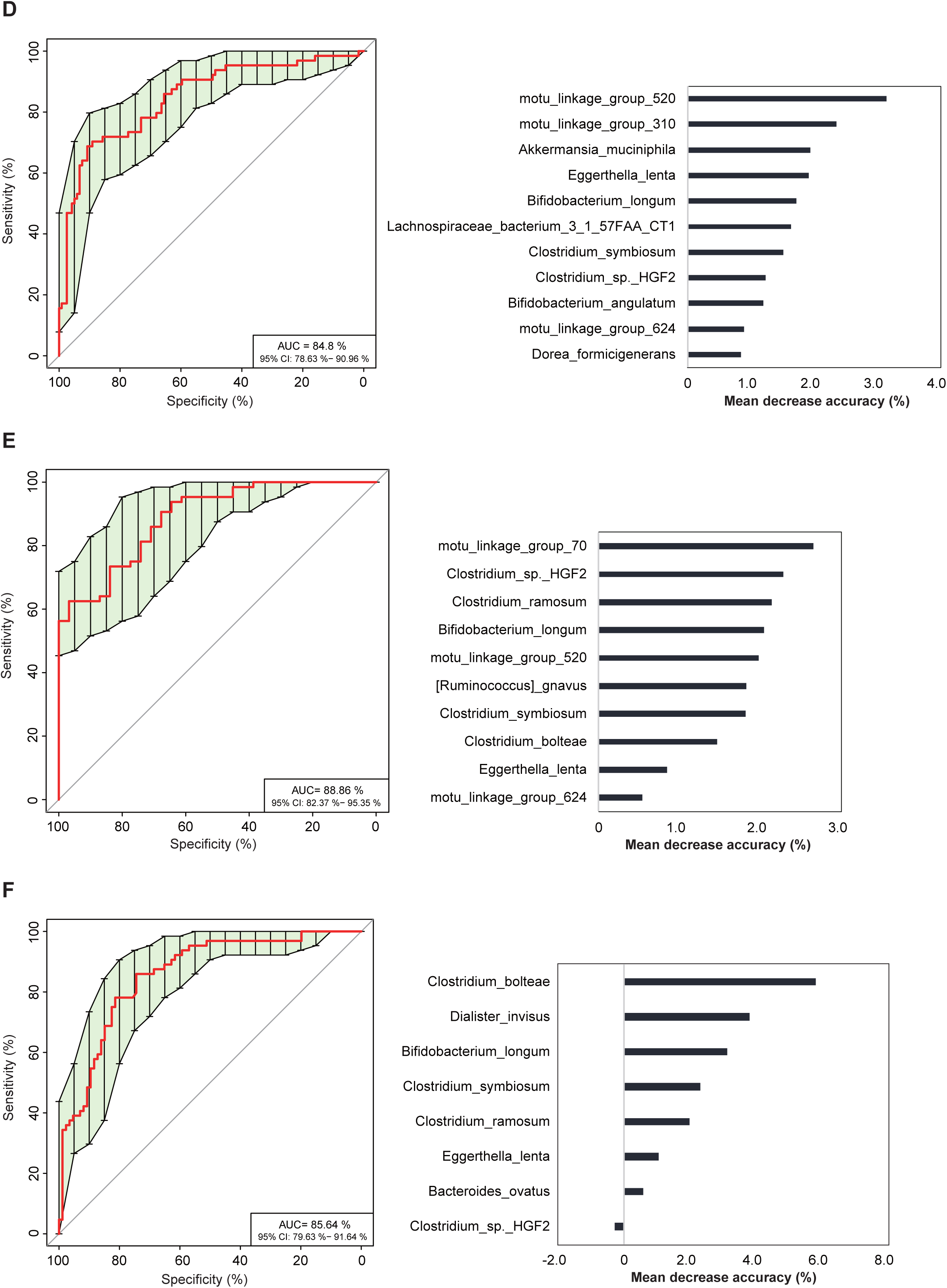
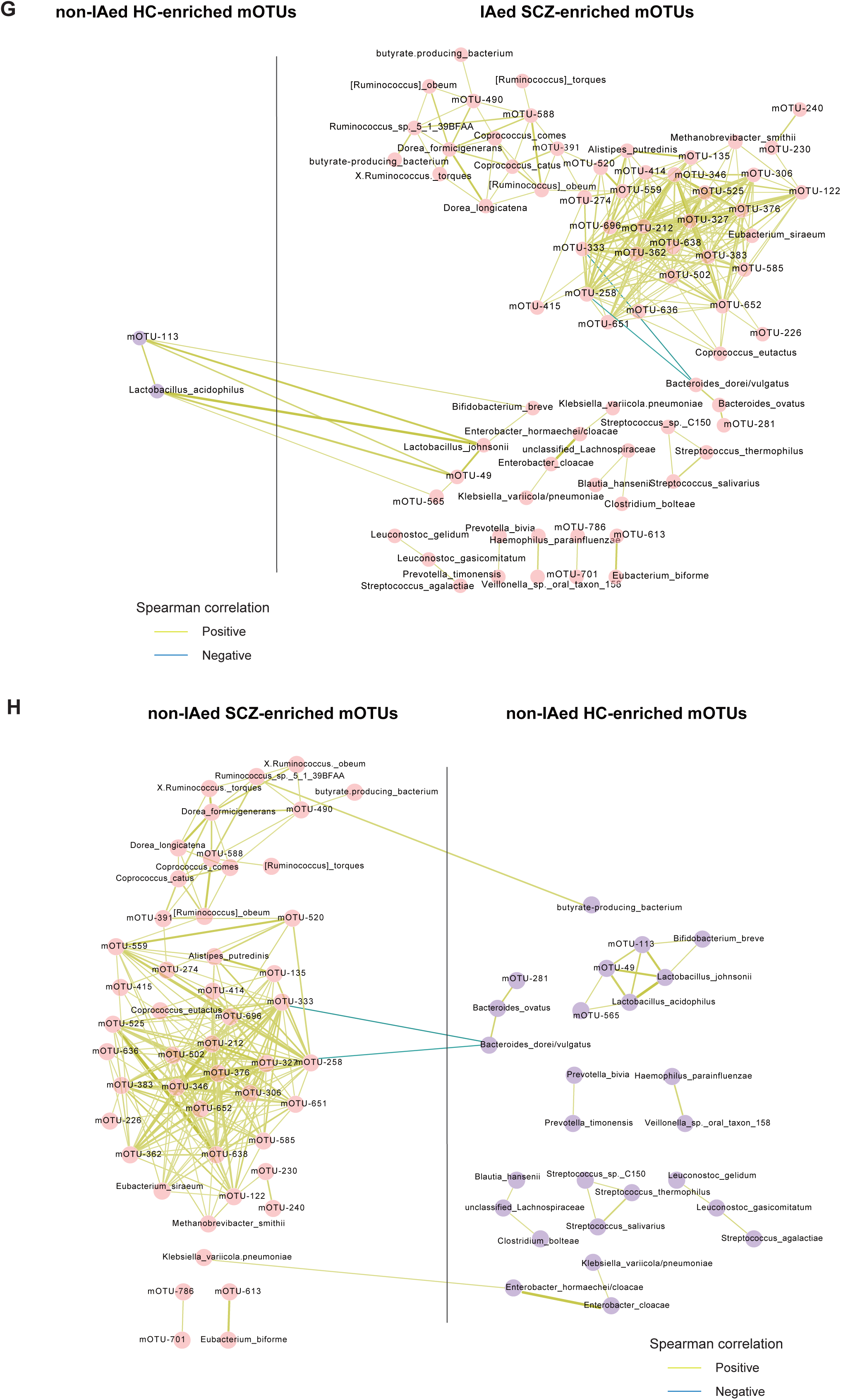
Gut microbiota alterations in immune-activated patients with schizophrenia. (A, B) α-diversity (Shannon index) and β-diversity (Bray-Curtis dissimilarity index of the gut microbiome). (C) Principal coordinates analysis on Bray-Curtis dissimilarity of mOTUs for immune-activated patients with schizophrenia, non-activated patients and non-activated controls. see panel legend for color key. (D-F) Receiver operating curves (ROC) according to 300 samples calculated by cross-validated random forest models. Area under ROC (AUCs) and the 95% confidence intervals (in parentheses) are also shown. (G, H) Network of mOTUs differentially enriched in healthy controls and schizophrenic patients. Node sizes reflect the mean abundance of significant mOTUs. mOTUs annotated to species are colored according to family (Red edges, Spearman’s rank correlation coefficient > 0.3, P < 0.05; blue edges, Spearman’s rank correlation coefficient <−0.3, *P* < 0.05;)

### Immune activation in schizophrenia is partially mediated through SCFA signaling

To address whether SCFA functionality mediates immune activation in schizophrenia, we explored the impacts of SCFAs on LPS-stimulated inflammatory response (IL-1β, IL-6, TNF-α secretion and NF-kB activation) of cultured PBMCs from each subgroup of participant defined by diagnosis and immune activation status (supplementary table S16). Pre-incubation of SCFAs significantly potentiated LPS-stimulated secretion of IL-1β, IL-6, TNF-α and NF-kB activation in the PBMCs from non-immune-activated patients, which was dependent on GPR41/43 signaling (all *P* < 0.01). The upregulated inflammatory response by SCFAs in PBMCs of non-immune-activated patients was not observed in immune-activated patients. For healthy controls, pre-incubation of SCFAs significantly attenuated LPS-induced IL-6 secretion and NF-kB activation in both immune-activated and non-immune-activated subgroups (all *P* < 0.01), which was independent of GPR41/43 signaling (all *P* > 0.05). In line with the diverse effects of SCFAs on immune response of PBMCs from the four different subgroups of participants, basal and LPS-induced cytokine responsiveness of PBMCs also varied among the subgroups. Both basal and LPS-stimulated inflammatory response in PBMCs were elevated in immune-activated patients compared with other three subgroups (all *P* < 0.01 figure 6A-C). Next, basal gene expression of two SCFA receptors and eight immune/inflammation mediators in PBMCs were quantified and compared among the four subgroups of participants. The immune-activated patients carried higher mRNA of *GPR41, GPR43, NF-kB, T-bet, RORC, STAT3, IL-1B, IL-6, IL-17a*, and *TNFA* and decreased *IL-10* and *Foxp3* in PBMCs, suggesting upregulated inflammatory response and enhanced differentiation potential into Th17 and Th1 effector cells, compared with the other three subgroups (all *P* < 0.001).

**Figure 6.**
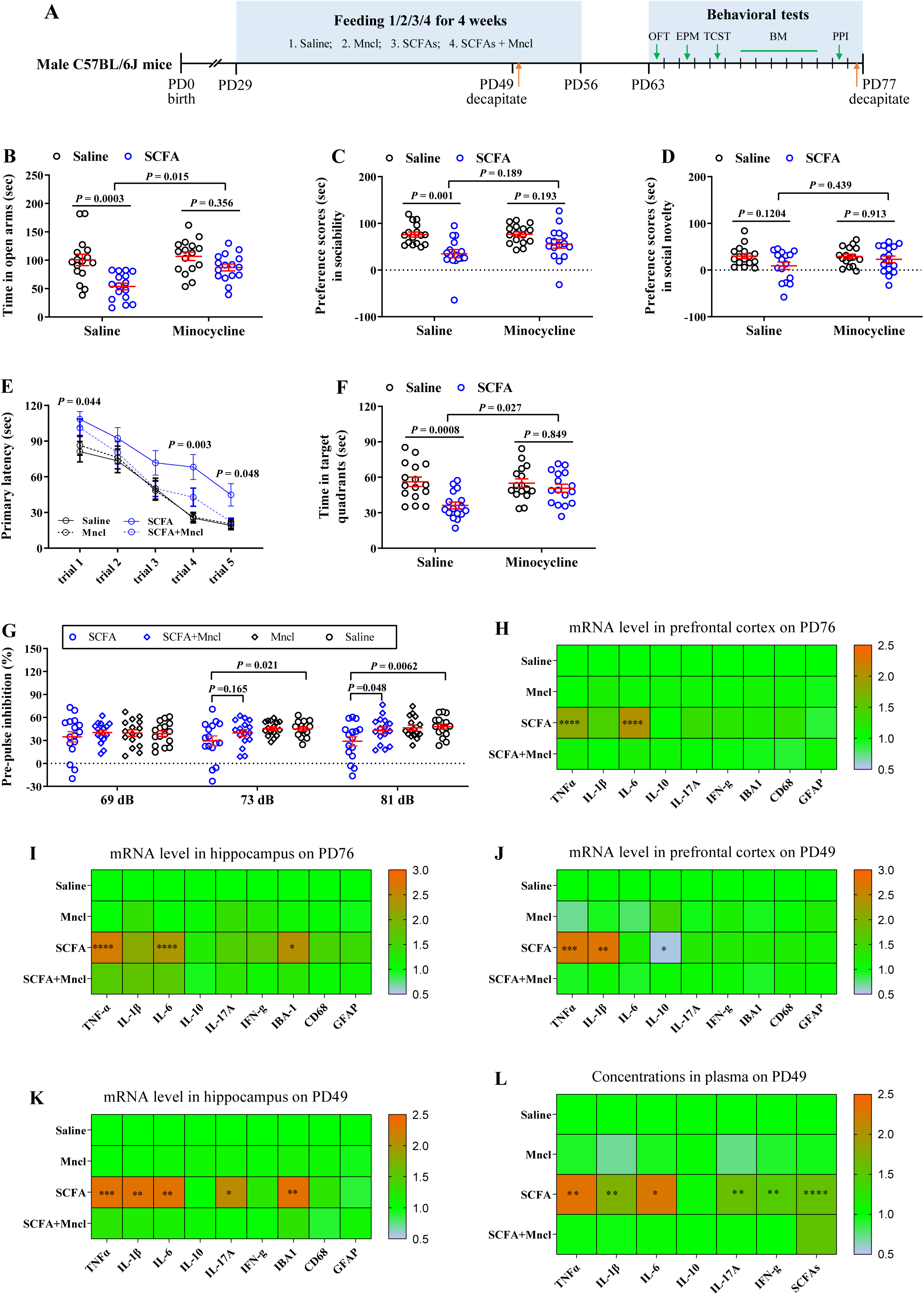
SCFAs contribute to schizophrenia-related immune activation in vitro cultured peripheral blood mononuclear cells.

Next, we investigated the impacts of serum small-molecule filtrates from different subgroups of participants on the secretion of pro-inflammatory cytokines and the gene expression profile of *GPR41, GPR43*, and 8 immune-related genes in cultured PBMCs from heathy volunteers. The small-molecule filtrate of serum from immune-activated patients and immune-activated controls significantly boosted inflammatory cytokines release by PBMCs compared to PBS exposure (all *P* < 0.01). SCFAs supplemented into the serum filtrates from immune-activated patients further elevated pro-inflammatory cytokine secretion, but SFCA supplementation into serum filtrate from immune-activated controls did not impact cytokine secretion. Antagonism of GPR41/43 weakened pro-inflammatory effects exerted by the serum filtrates of immune-activated patients on PBMC (all *P* < 0.01), but did not significantly impact the serum filtrates from immune-activated controls (all *P* > 0.01). The effects of SCFA supplementation on cytokine production were attenuated by blocking GPR41/43 (all *P* < 0.01). In line with SCFAs in serum, SCFAs in small-molecule filtrate of serum were also increased in immune-activated patients compared to the other three subgroups. Small-molecule filtrate of serum from immune-activated patients significantly increased gene expression of *GPR41, GPR43, NF-kB, T-bet, RORC, STAT3, IL-6* and decreased mRNA of *IL-10* and *Foxp3* in cultured PBMCs of healthy volunteers, which is similar to the expression signature in their basal PBMCs cultured *in vitro*.

### Chronic elevation of SCFAs in peri-puberty induced schizophrenia-related abnormal behaviors in adult mice

To address the role of SCFAs in schizophrenia, we investigated the impacts of SCFA feeding on immune function and behaviors of mice (Experiments 1, 2 in figure 1). Four weeks of excessive SCFA ingestion in adult mice did not induce marked behavioral changes and inflammatory response in blood or gut (online supplementary figure S1). However, excessive intake of SCFAs in peri-puberty, an important period of neurodevelopment[46], induced schizophrenia-relevant abnormal behaviors, including elevated anxiety level, impaired social behaviors, spatial memory and PPI in adulthood (figures 6A-G). Moreover, feeding SCFAs in peri-puberty increased pro-inflammatory cytokines (TNF-α and IL-6) in PFC and hippocampus in adulthood (i.e. 20 days after stopping drinking SCFAs; figure 6H, I), as well as pro-inflammatory responses in both blood and brain and elevated serum SCFAs on PD49 (i.e. had drunk SCFAs for three weeks; figure 6J, K, L, M), compared with saline-treated mice (online supplementary figure S2). Moreover, oral treatment of SCFA-fed animals with the anti-inflammatory compound minocycline attenuated or eliminated the SCFA-induced abnormal behaviors and systemic immune activation (all *P* < 0.01, figures 6A-G). Feeding SCFA in peri-puberty did not exert significant effects on locomotor, inflammatory level in blood and serum SCFAs and weight in adult mice (online supplementary figure S2).

### SCFAs exacerbate immune activation and expression of schizophrenia-related behaviors in a mouse model of schizophrenia

To recapitulate the pathophysiology underlying SCFA elevation in immune-activated patients with schizophrenia, we increased circulating SCFAs of adult mice with MIA (mouse immune activation) via four-weeks of SCFA supplementation in drinking water. And then we investigated the effects of SCFA elevations on immune status and behavior symptoms in a MIA model of schizophrenia (figure 7A).

**Figure 7.** SCFAs contribute to schizophrenia-related behavioral symptoms and immune activation in mice.

**Figure 8.** SCFAs enhance the expression of schizophrenia-related behavioral symptoms in mice model of maternal immune activation.

Although SCFA intake in adult mice did not significantly affect immunity and behavior, excessive SCFA intake potentiated MIA-induced schizophrenia-relevant behaviors, including more profoundly impaired PPI and social behavior (all *P* < 0.01, figure 7B-C). Moreover, SCFA intake also upregulated several pro-inflammatory cytokines in serum (IL-6, TNF-α), PFC (IL-6), and hippocampus (IL-6) of adult offspring of MIA mice (all *P* < 0.01, figure 7D and Supplementary table S17). Again, minocycline significantly decreased the effects of SCFAs on expression of schizophrenia-relevant behaviors and pro-inflammatory cytokines in adult offspring of MIA mice (all *P* < 0.01, figures 7D). Feeding SCFA in drinking water significantly increased cecum SCFA and circulating SCFA in mice (online supplementary table S18). Moreover, prenatal immune activation potentiated SCFA elevation during chronic SCFA intake, which is accompanied by increased gut permeability in mice (figures 7E, F).

## DISCUSSION

In the present study, we elucidate causative roles of gut microbiota dysbiosis-induced SCFA elevation in schizophrenia pathology, especially for immune activation and cognitive impairment. This conclusion is based on two lines of evidence from human and animal, respectively. Although SCFAs are generally regarded as beneficial microbial metabolites for human health, our results, together with several lines of evidence [47, 48, 49, 50] indicate that, like a double-edge sword, excessive SCFA beyond physiological concentration can interfere with biological functioning of host. To the best of our knowledge, this is the first study to show that higher fecal SCFA excretion and circulating SCFA are presented in schizophrenia. Generally, circulating SCFAs may better represent SCFA production and absorption[50]. Increased SCFA in both circulation and feces of patients with schizophrenia suggests that SCFA production by microbiota is too excessive in gut to be decreased to normal level by increasing absorption. In line with this speculation, several SCFA-producing bacterial species are more abundant in gut of patients and display moderate correlations between serum and fecal SCFAs. To more precisely evaluate the status of bacterial production and/or intestinal absorption of SCFA in schizophrenia, SCFA release in different locations along the intestine and its concentration in portal vein, hepatic vein and mesenteric vein[51] should be measured in future human study.

Elevated SCFA are also presented in autism spectrum disorder[47], periodontitis[52], obesity, hypertension and cardiometabolic disease[9]. Shared pathological pathways/systems exist between schizophrenia and these disorders, including gut dysbiosis, perinatal inflammation[53], and systemic immune activation/chronic low-grade inflammation[54]. These overlapped pathways imply that, apart from dysbiotic gut microbiota, dysfunctional immune system may also underpin elevated SCFA production and/or absorption. Previous studies propose that increased SCFA in the systemic circulation may be facilitated by increased intestinal permeability [39, 55, 56], which are apparent in some patients with schizophrenia [57, 58]. Our human data reveal that sCD14, serological surrogate markers for both bacterial translocation and innate immune activation, are upregulated in schizophrenia and positively correlated with circulating SCFAs. In addition, up-regulated SCFA in schizophrenia is mainly presented in immune-activated patients. These findings suggest potential mechanistical links between leaky gut, systemic immune activation, and excessive entering of SCFA into circulation. Consistent with these findings in human, animal data demonstrate that prenatal immune activation via viral mimetic poly(I:C) challenge to pregnant maternal mice induces defective gut barrier in adult offspring mice and in parallel potentiates oral SCFAs intake-induced upregulation of intestinal expression of SCFA receptors and serum SCFA. The concentrations of SCFAs in colon lumen and intestine wall at basal condition and after three-week of SCFAs drinking are similar between the mice with prenatal immune activation and control mice. These results indicate that intrinsic immune dysfunction and leaky gut promotes to entering of SCFA into blood when excessive SCFAs are presented in gut. Combining evidence from human and mice, we propose that both increased gut permeability derived from dysfunctional immune systems and increased formation of exogenous SCFA in gut derived from dysbiotic microbiota contribute to increased SCFAs in circulation in schizophrenia.

Circulating SCFAs upregulation are not only contributed by immune dysfunction and leaky gut, but also further contribute to immune activation. It is like a positive feedback to exacerbate the imbalanced immune response in schizophrenia. The contributory effects of SCFAs on immune activation are also revealed by both human study and mice study. Identifying systemic effects of SCFAs on host physiology are important for understanding their roles in disease pathophysiology. Our human data indicate that excessive SCFA secretion in blood is positively associated with systemic immune activation and cognitive impairment, but not with severity of psychotic symptom in schizophrenic patients. Clinical investigation indicate that cognitive impairment is intrinsic deficit of schizophrenia and largely resistant to current treatment. Similarly, our data revealed that three-month antipsychotics treatment, although improve cognitive symptoms to some extent, their cognitive function remained significantly impaired relative to controls. Moreover, treatment did not significantly affect serum SCFA level, abundance of SCFA-producing bacteria and the proportion of immune activation in patients compared with those at baseline. Such stable alterations highlight a unique axis of dysbiotic gut microbiota-excessive SCFA-immune activation-cognitive impairment in schizophrenia. Previous studies have identified many pathways mediating immune-brain communication[59, 60]. Increasing evidence suggest that dysregulated peripheral inflammatory signaling may increase astrocyte tryptophan-dioxygenase activity in the brain and subsequently disturb glutamate transmission and cholinergic signaling, which are in part responsible for negative and cognitive symptoms of schizophrenia[61, 62]. Our human data reveal serum SCFA acts as an upstream mediator linking gut microbiota dysbiosis and dysregulated immune activation. Serum SCFAs more accurately reflect intestinal SCFAs production than fecal SCFA [63, 64] and can be distributed throughout the body to regulate physiology of remote organ[65]. Most absorbed butyrate is metabolized in intestinal epithelial cells, while acetate and propionate reaches the systemic circulation at higher concentrations[66]. Similarly, our data reveal butyrate is not elevated in serum of patients despite increased concentration in feces. So, we propose that increased acetate and propionate in serum exerts systemic effects on immune dysfunction in schizophrenia.

Our human classification based on four immune mediators in serum reveals ∼45% patients and ∼15% healthy controls display marked immune activation status compared with other participants. These results are consistent with two previous studies reporting that immune dysfunction is presented in a subgroup of patients with schizophrenia but not all patients [41, 67], despite different immune/inflammatory markers are selected by three studies. Serum SCFA is upregulated in immune-activated patients but not in immune-activated healthy controls. And, gut microbiota analysis also identifies unique bacterial signature underpinning immune-activated schizophrenia that can distinguish these patients from others. These results highlight intrinsic factors driving immune activation in schizophrenia is naturally different with those in general population. Some immune-related risk factors for schizophrenia, such as genetic variations in immune-related genes and early-life adversity and immune activation, may confer them biological heterogeneity in the response of immune system and gut microbiota to exogenous stimulation[68, 69].

Another important finding in the present study is that excessive SCFA level contributes to schizophrenia-related immune activation revealed by analyses in both *in-vitro* human immune cell and mouse model of schizophrenia. The cellular targets of SCFA action to modulate immune-related cell function, cellular signaling and gene expression are very various, and include histone deacetylase activity, activation of free fatty acid G-coupled receptor and mitochondrial inflammatory signaling cascades, which may or may not be mutually reinforcing[70]. SCFAs promote T cell differentiation into both effector and regulatory T cells to promote either immunity or immune tolerance depending on immunological milieu[18]. On one hand, SCFAs, particularly butyrate, are known to suppress intestinal inflammation[71, 72] and down-regulate nitric oxide and proinflammatory cytokines in multiple cells or tissues[73, 74]. On the other hand, over-stimulation of SCFAs also up-regulate inflammatory cytokines in the colon of pig[75] and induce ureteritis and hydronephrosis in mice[17]. Increasing evidence indicate that elaborate regulation of SCFA on immune system are dose- and tissue-dependent, and exert di□erent e□ects at key developmental time periods[76]. The effects of SCFA on immune response of PBMC are dependent on diagnosis and intrinsic immune status of donors. SCFA exerts opposite effects (enhancement vs attenuation) LPS-induced pro-inflammatory cytokine release in the PBMCs from non-immune-activated patients and from healthy controls, respectively. These results again highlight intrinsic vulnerability factors for immune in schizophrenia patients differ from controls. Significantly different expression signature of immune-related genes in PBMCs among four subgroups of participants may explain the differences in SCFA effects between four subgroups. The PBMCs from immune-activated patients have exposed to excessive SCFA levels in patients’ serum and display increased monocyte proportion, elevated expression of pro-inflammatory cytokines, increased Th1 and Th17 differentiation on baseline, compared with other 3 subgroups. The cellular signaling of these PBMCs targeted by SCFA may be saturated by pre-exposure of SCFA in serum, which disables them to act to *in-vitro* SCFA stimulation. In non-immune-activated patients, *in-vitro* SCFA potentiates LPS-induced inflammatory response of PBMC and these effects can be blocked by inhibitor of G0/i. Moreover, small-molecule filtrate of serum from immune-activated patients enhance LPS-stimulated inflammatory response of normal PBMC, suggesting small molecules in serum of these patients are driving factor for immune activation. Compared with immune-activated controls, small-molecule filtrate of serum from immune-activated patients includes higher SCFA and also exert more powerful effect on inflammatory mediator secretion of PBMC. Mechanistically, the effects of small-molecule filtrate of serum from immune-activated patients on immune response of PBMC can be attenuated by inhibition of SCFA signaling. These data demonstrate that cellular signaling activated by enteric microbial-derived SCFA partially mediates the contributory effects of serum metabolites of patients on immune activation. It is notable that SCFA signaling only account for a fraction of effects of serum metabolites, more studies in future need to be done to identify more effector metabolites for immune activation.

Our animal study provides more detailed mechanisms underlying the roles of SCFAs in immune activation and aberrant brain function in schizophrenia. Mouse model of schizophrenia revealed a mechanistic link between microbial-derived SCFAs (acetate and propionate) and immune activation and cognitive impairments. Our data demonstrate excessive SCFA intake in adolescent *per se* induced some schizophrenia-relevant behaviors, including elevated anxiety level and impaired spatial memory and social behaviors in mice. Although the mechanisms by which SCFA contribute to behavioral abnormality in mice is still unclear, some neural pathways affected directly SCFA should be considered, including adverse metabolic effects, neuroinflammation, microglia activation, electrophysiological disturbances as well as disruptions in lipid, mitochondrial and redox metabolism[26, 37, 77, 78]. Consistent with previous studies, prenatal immune stimulation by poly I:C induced schizophrenia-like behaviors in adult, including deficit in learning and memory[79]. Moreover, our data revealed significant interaction between prenatal immune stimulation and excessive SCFA intake in adolescent on systemic immune activation and the expression of impaired memory and social behaviors in mice. The parallel increase in serum SCFA and pro-inflammatory cytokines, neuroinflammation marker expression in cortex, and neurochemical molecules in hippocampus, and memory impairment. Moreover, oral treatment of SCFA-treated mice with the anti-inflammatory agent minocycline partially reduce serum pro-inflammatory response and alleviated neuroinflammation and improve behavioral impairments. These findings demonstrate that SCFA induces cascade changes in these pathways to enhance behavioral impairment caused by prenatal immune activation. The systemic immune activation are key pathways influenced by SCFA to modulate abnormal behaviors expression in adult of prenatally immune-activated mice. Previous studies have revealed mechanisms by which peripheral immune activation induce neuroinflammation and psychopathology. Our study identified gut microbiota dysbiosis-induce excessive SCFA production is upstream mediators for immune activation in schizophrenia. In summary, findings from human and mouse study identify increased SCFA production resulting from dysbiotic gut microbiota and subsequent potentiate systemic immune activation and contribute to cognitive impairment. The SCFA-producing bacteria-serum SCFA-immune activation can be possible therapeutic targets for schizophrenia.

## Acknowledgements

This study was supported by the Clinical Research Award of the First Affiliated Hospital of Xi’an Jiaotong University (No. XJTU1AF-CRF-2016-005), Innovation Team Project of Natural Science Fund of Shaanxi Province (2017KCT-20), and Key Program of Natural Science Fund of Shaanxi Province (2018ZDXD-SF-036).

